# More comprehensive proprioceptive stimulation of the hand amplifies its cortical processing

**DOI:** 10.1101/2021.02.24.432547

**Authors:** Maria Hakonen, Timo Nurmi, Jaakko Vallinoja, Julia Jaatela, Harri Piitulainen

## Abstract

Corticokinematic coherence (CKC) quantifies the phase coupling between limb kinematics and cortical neurophysiological signals reflecting proprioceptive feedback to the primary sensorimotor (SM1) cortex. We studied CKC to proprioceptive stimulation (*i*.*e*. movement-actuator-evoked movements) of right-hand digits (index, middle, ring and little) performed simultaneously or separately. CKC was computed between magnetoencephalography (MEG) and finger acceleration signals. The strongest CKC was obtained by stimulating the fingers simultaneously at fixed 3-Hz frequency, and can, therefore, be recommended as design for fast functional localization of the hand area in the primary sensorimotor (SM1) cortex using MEG. The peaks of CKC sources were concentrated in the hand region of the SM1 cortex, but did not follow consistent somatotopic order. This result suggests that spatial specificity of MEG is not sufficient to separate proprioceptive finger representations of the same hand adequately or that their representations are overlapping.

## Introduction

Corticokinematic coherence (CKC) quantifies the phase coupling between limb kinematics (e.g. hand acceleration or contractile force, Piitulainen et al., 2013a) and cortical neurophysiological signals measured with magnetoencephalography (MEG, Piitulainen et al., 2013b) or electroencephalography (EEG) in adults (Piitulainen et al., 2020) and even in infants (Smeds et al., 2017). CKC peaks at the movement frequency and its harmonics in the primary sensorimotor cortex (SM1) contralateral to the moving limb (Bourguignon et al., 2011; Jerbi et al., 2007). CKC is strong for repetitive finger (Piitulainen et al., 2013b), toe (Piitulainen et al., 2015) and ankle (Piitulainen et al., 2018b) movements, and follows the respective somatotopic cortical representations. Thus, CKC is a robust tool to pinpoint, e.g., functional hand representation (Bourguignon et al., 2013b), that can be valuable information when planning a brain surgery. CKC has shown to activate the SM1 cortex similarly in different rates (∼1–12 Hz) of voluntary (Marty et al., 2015) and movement-actuator induced hand movements (Piitulainen et al., 2015), with no differences in the CKC strength or source location.

CKC primarily reflects somatosensory afference to the SM1 cortex (Bourguignon et al., 2015; Piitulainen et al., 2013b), since active (volitional) and passive (evoked by an investigator) movements elicited similar strength and cortical location of CKC. Moreover, CKC is not affected by the level of tactile contamination, and the cortical CKC source is spatially distinct from the tactile source in of the same finger (Piitulainen et al., 2013b). These observations suggest that CKC primarily reflects proprioceptive afference (presumably from the muscle spindles) to the SM1 cortex arising from the rhythmic movement (Bourguignon et al., 2015; Piitulainen et al., 2013b). Thus, CKC can be used to quantify degree or extent of cortical proprioceptive processing, and may be utilized to identify impairments in proprioceptive pathways in various motor disorders (e.g. Marty et al., 2019) or in healthy aging (Piitulainen et al., 2018b). Finally, CKC has shown to be a reproducible tool to follow cortical proprioception at group level both for MEG and EEG (Piitulainen et al., 2018a, 2020).

The primary aim of this study was to examine whether the CKC strength or cortical source location differs between proprioceptive stimulation (*i*.*e*. movement-actuator-evoked movements) of the right-hand digits (D2–D5: index, middle, ring and little). We aimed to determine whether a comprehensive multi-finger stimulation would improve robustness and time efficiency of CKC based functional localization of the hand SM1 cortex using MEG. Three conditions were tested: (1) simultaneous stimulation of all four fingers at 3-Hz frequency (*simultaneous* _constant-*f*_), (2) stimulation of each finger separately at 3-Hz frequency (*separate*) and (3) simultaneous stimulation of the fingers at finger-specific frequencies (at 2, 2.5, 3 and 3.5 Hz, *simultaneous* _varied-*f*_).

We had four hypotheses. The first hypothesis (H1) was that the simultaneous stimulation of the four fingers would result in stronger CKC due to stronger proprioceptive afference to the SM1 cortex compared to the separate-finger stimulation. The second hypothesis (H2) was that the strength of CKC is similar both for *simultaneous* _varied-*f*_ and *separate* conditions, because the stimulation of the finger at a finger-specific frequency is analogous to stimulating the finger separately. The benefit here is that *simultaneous* _varied-*f*_ approach would provide more time efficient CKC recording if cortical representations or CKC values of individual fingers are of interest (*e*.*g*. in specific clinical conditions). The third hypothesis (H3) was that the most dexterous index finger, with presumably larger cortical proprioceptive representation in the SM1 cortex, would show the strongest CKC, and the least dexterous ring finger the weakest CKC (for studies comparing finger dexterity, see Aoki et al., 2003; Häger-Ross and Schieber, 2000; Ingram et al., 2008; Kinoshita et al., 1996; Reilly and Hammond, 2000; Swanson et al., 1974; Zatsiorsky et al., 1998). Finally, the fourth hypothesis (H4) was that the cortical source location does not vary significantly between fingers in our healthy participants, and thus each finger representation would similarly represent the hand region of the SM1 cortex when assessed with MEG.

## Results

Figure 1 shows the movement actuator as well as averaged MEG and acceleration signals measured during proprioceptive stimulation for a representative participant in three experimental conditions. We included only successful recordings of participants with clear CKC topographies and statistically significant (p < 0.05) CKC peaking at the stimulation frequency in the final analysis (n = 16–18 participants depending on the condition). The assumptions of normality or sphericity were not violated in the data. As expected, the results from sensor and source level analyzes were replicated well at the individual level. There were no systematic between-finger or between-condition differences in the MEG gradiometer (sensor) pair in which CKC peaked. CKC peaked in MEG422-MEG423 gradiometer pair (50% of the cases) or in a gradiometer pair just adjacent to it.

**Figure 1.**
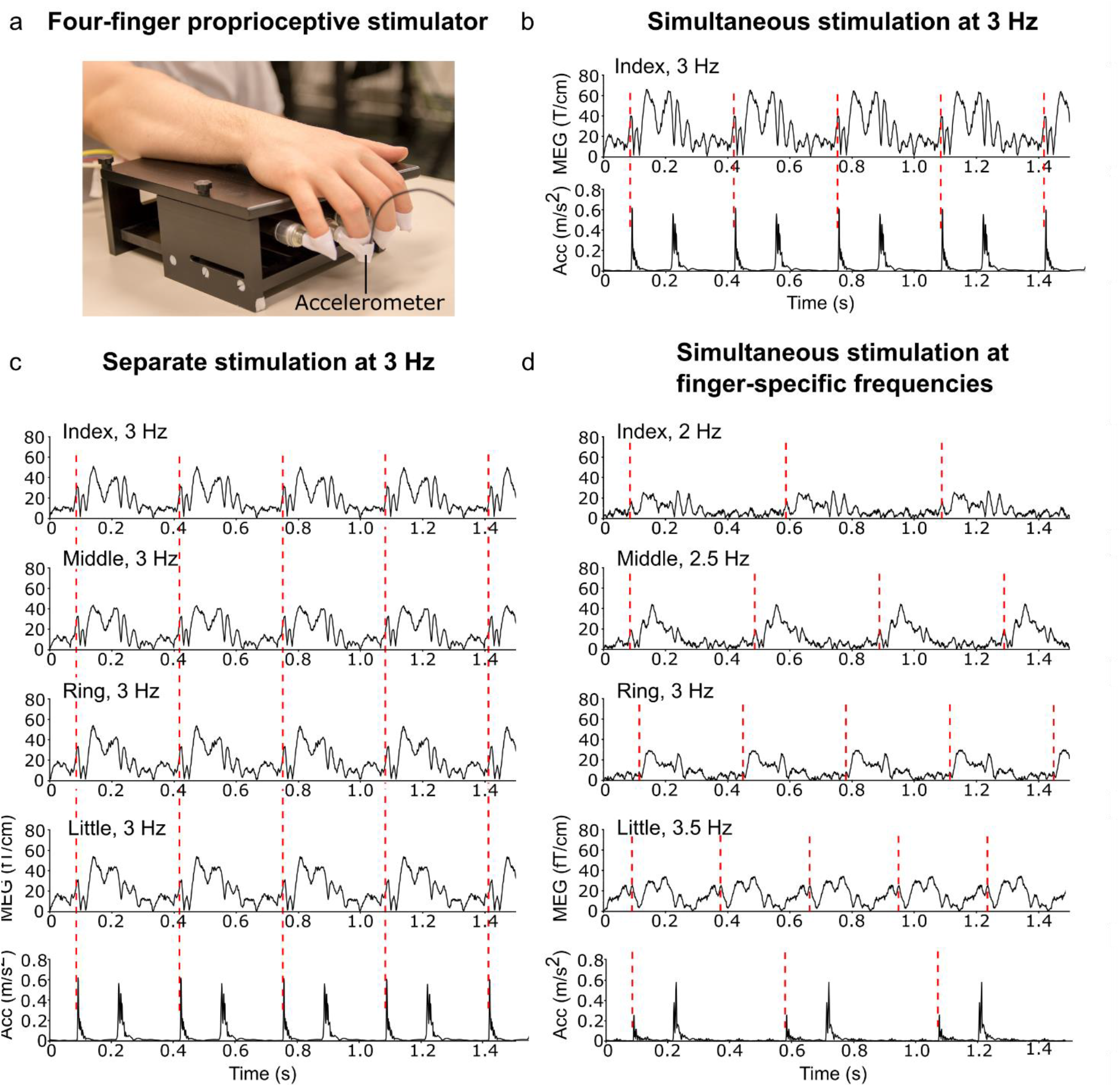
Proprioceptive stimulator, sustained-MEG fields for each finger and acceleration magnitude for the index finger. (a) The four-finger proprioceptive stimulator. Please, note that the figure is only for visualization purposes and does not include all four accelerometers. (b–d) Averaged MEG responses (vector sum of the peak gradiometer pair) for each finger and acceleration magnitude (Euclidean norm of the three orthogonal components) for the index finger in all three conditions. The red dashed line indicates an onset of the flexion phase of the continuous flexion-extension movement.

### Stronger CKC to simultaneous than separate-finger stimulation at 3 Hz (H1)

The total number of accepted trials did not differ significantly between the conditions (*simultaneous* _constant-*f*_ : 417 ± 32, *separate*: 419 ± 23, two-sample t-test, p = 0.79, n = 18). **Figure 1a** and **Table 1** present CKC strength for simultaneous and separate 3-Hz stimulation both at sensor and source levels. In line with our first hypothesis, CKC was stronger when the fingers were stimulated simultaneously than when they were stimulated separately, presumably reflecting stronger proprioceptive afference to the SM1 cortex. This effect was detected for all fingers.

**Table 1.** CKC strength for separate and simultaneous 3-Hz stimulations at sensor and source levels.

| H1 (N=18) | separate | simultaneous<sub>constant</sub> | p | F | df1 | df2 |
| --- | --- | --- | --- | --- | --- | --- |
| **Sensor level** |  |  |  |  |  |  |
| Finger average | $0.42 \pm 0.03$ | $0.69 \pm 0.03$ | **<0.001** | 191.03 | 1 | 17 |
| *Interaction* | - | - | 0.05 | 2.76 | 3 | 51 |
| *Index* | $0.40 \pm 0.04$ | $0.69 \pm 0.04$ | **<0.001** | - | - | - |
| *Middle* | $0.37 \pm 0.04$ | $0.69 \pm 0.03$ | **<0.001** | - | - | - |
| *Ring* | $0.46 \pm 0.04$ | $0.69 \pm 0.03$ | **<0.001** | - | - | - |
| *Little* | $0.46 \pm 0.04$ | $0.69 \pm 0.03$ | **<0.001** | - | - | - |
| **Source level** |  |  |  |  |  |  |
| Finger average | $0.39 \pm 0.02$ | $0.50 \pm 0.02$ | **<0.001** | 19.56 | 1 | 17 |
| *Interaction* | - | - | **<0.01** | 4.47 | 3 | 51 |
| *Index* | $0.34 \pm 0.03$ | $0.50 \pm 0.03$ | **<0.01** | - | - | - |
| *Middle* | $0.36 \pm 0.03$ | $0.50 \pm 0.03$ | **<0.001** | - | - | - |
| *Ring* | $0.40 \pm 0.03$ | $0.50 \pm 0.03$ | **<0.001** | - | - | - |
| *Little* | $0.43 \pm 0.03$ | $0.50 \pm 0.03$ | **<0.001** | - | - | - |

### Weaker CKC to simultaneous stimulation at finger-specific frequencies than separate stimulation at 3 Hz (H2)

The total number of accepted trials did not differ significantly between the conditions (*simultaneous* _varied-*f*_ : 419 ± 23 trials and *separate:* 419 ± 23, n = 16). In contrast to our second hypothesis, CKC was weaker when the fingers were stimulated simultaneously at the finger-specific frequencies (**Fig. 2b, Table 2**), indicating that the simultaneous approach is not analogous with the separate finger stimulation. Furthermore, the reductions in CKC strength were finger-specific (**Fig. 2c, Table 2**). CKC was weaker for the index, ring and little fingers, but not for the middle finger in the *simultaneous* _varied-*f*_ condition. The results were similar at the source level with exceptions that CKC was weaker also for the middle finger, and no difference was found between the conditions for the index finger.

**Table 2.** CKC strength for separate 3-Hz stimulation and simultaneous stimulation at finger-specific frequencies.

| H2 (N=16) | separate | simultaneous<sub>varied</sub> | p | F | df1 | df2 |
| --- | --- | --- | --- | --- | --- | --- |
| **Sensor level** |  |  |  |  |  |  |
| Finger average | 0.43 ± 0.04 | 0.30 ± 0.02 | <**0.001** | 24.91 | 1 | 15 |
| *Interaction* | - | - | <**0.001** | 6.45 | 3 | 45 |
| *Index* | 0.39 ± 0.05 | 0.31 ± 0.04 | <**0.05** | - | - | - |
| *Middle* | 0.38 ± 0.04 | 0.33 ± 0.03 | 0.054 | - | - | - |
| *Ring* | 0.47 ± 0.04 | 0.32 ± 0.03 | <**0.001** | - | - | - |
| *Little* | 0.46 ± 0.05 | 0.25 ± 0.03 | <**0.001** | - | - | - |
| **Source level** |  |  |  |  |  |  |
| Finger average | 0.38 ± 0.02 | 0.25 ± 0.02 | <**0.001** | 66.09 | 1 | 15 |
| *Interaction* | - | - | <**0.001** | 19.56 | 3 | 45 |
| *Index* | 0.34 ± 0.03 | 0.31 ± 0.03 | 0.23 | - | - | - |
| *Middle* | 0.36 ± 0.03 | 0.23 ± 0.03 | <**0.001** | - | - | - |
| *Ring* | 0.41 ± 0.03 | 0.31 ± 0.02 | <**0.001** | - | - | - |
| *Little* | 0.43 ± 0.03 | 0.16 ± 0.02 | <**0.001** | - | - | - |

**Figure 2.**
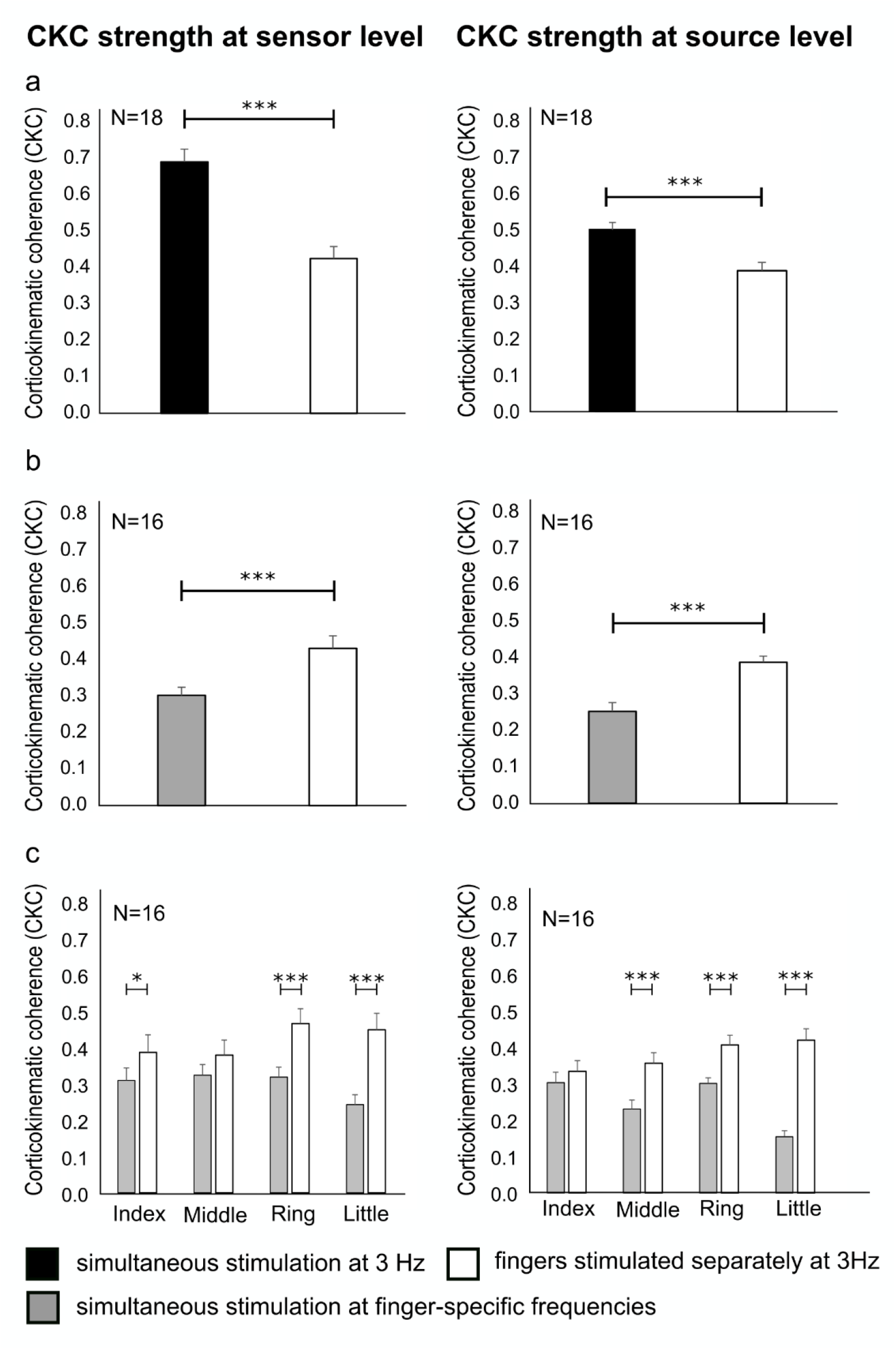
CKC strength for all conditions at the sensor and source levels. Average CKC strength for (a) simultaneous *versus* separate stimulation at 3 Hz, (b) simultaneous stimulation at the finger-specific frequencies *versus* separate stimulation at 3 Hz, and (c) individual fingers when stimulated simultaneously at the finger-specific frequencies and separately. * p < 0.05, ** p < 0.01, *** p < 0.001.

### CKC strength reflects finger dexterity and functional dominance (H3)

Figure 3. shows the CKC strength for individual fingers elicited by separate stimulation and by simultaneous stimulation at the finger-specific frequencies (n = 18). In contrast to our third hypothesis, the most dexterous index finger did not show the strongest CKC when the fingers were stimulated separately. Instead, it seems that the dexterity of the finger decreases CKC. CKC was stronger for the ring (0.46 ± 0.04) and little (0.46 ± 0.04) fingers when compared to the middle finger (0.37 ± 0.04, p < 0.05, **Fig. 3a**). Similar results were observed at the source-level, where the CKC was stronger for the ring (0.41 ± 0.03, p < 0.01) and little (0.43 ± 0.03, p < 0.001) fingers when compared to the middle finger (0.36 ± 0.03, **Fig. 3a**). In addition, at the source-level, CKC was stronger also for the ring (p < 0.05) and little (p < 0.01) fingers when compared to the index finger (0.34 ± 0.03).

**Figure 3.**
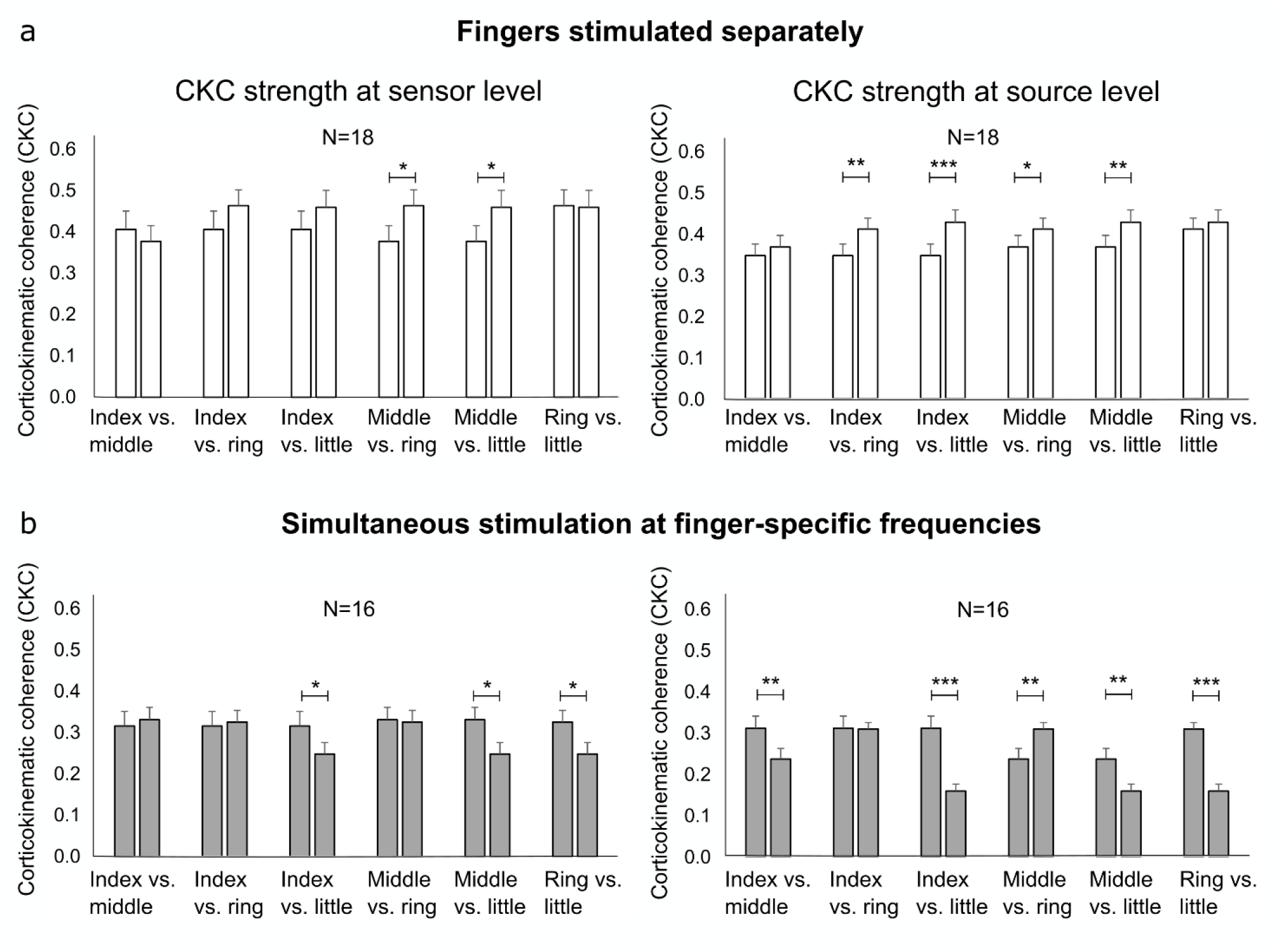
CKC strength for individual fingers. Average CKC for (a) the separate stimulation of individual fingers and (b) for the simultaneous stimulation at the fingers at their specific frequencies. * p < 0.05, ** p < 0.01, ***p < 0.001.

When the fingers were stimulated simultaneously at the finger-specific frequencies, the results were partly in line with our third hypothesis. The more dexterous fingers showed stronger CKC than the least dexterous little finger. CKC was stronger for the index (0.31 ± 0.04, p < 0.02, 2 Hz), middle (0.33 ± 0.03, p < 0.03, 2.5 Hz) and ring (0.32 ± 0.02, p < 0.02, 3 Hz) fingers than for the little finger (0.25 ± 0.03, 3.5 Hz, **Fig. 3b**). Again, the source level results were similar: CKC was stronger for the index (0.31 ± 0.03, p < 0.001), middle (0.23 ± 0.03, p < 0.004) and ring (0.31 ± 0.02, p < 0.001) fingers than for the little finger (0.16 ± 0.02, **Fig. 3b**). Moreover, CKC was stronger for the index (p < 0.003) and ring (p < 0.002) fingers than for the middle finger. In this condition, the CKC strength likely partly reflects the dominance or contribution of a given finger to the overall hand proprioceptive processing in the SM1 cortex.

### CKC source locations were concentrated on hand region of the SM1 cortex (H4)

An average between-finger distance was 8.0 ± 2.8 mm (N = 16) across the finger source locations obtained by separate stimulation and simultaneous stimulation at finger specific frequencies. Sixteen participants were included in all comparisons between source locations. In contrast to our fourth hypothesis, the source locations of the fingers were partly distinct, but did not follow the consistent somatotopic pattern (**Fig. 4, Table 3**) indicated by Penfield’s homunculus (Penfield and Boldery, 1937; Penfield and Rasmussen, 1950) and subsequent studies (Hari et al., 1993; Nakamura et al., 1998). The somatosensory representation of the index finger has shown to be the most dorsal and inferior one along the central sulcus followed by the representations of the middle and ring fingers and, finally, the most ventral and superior representation of the little finger. Nevertheless, there were significant differences between the proprioceptive representations of the fingers. The CKC peak location was more medial for the ring (3.4 mm, p < 0.03) and little (4.2 mm, p < 0.01) fingers than for the index finger (x = **–**47.6 ± 1.1 mm) when the fingers were stimulated separately. Additionally, the ring finger was 3.9 mm more superior than the index finger (z = 54.5 ± 1.7 mm, p < 0.04). According to a visual inspection of the participants’ CKC source locations, the index and little fingers roughly followed the somatotopic arrangement in 6 out of 16 participants, the index finger being represented more dorsal and inferior to the little finger.

**Table 3.** The grand average MNI coordinates of CKC peak source locations.

| Finger | simultaneous<sub>constant-f</sub>, (N=18) |  |  | separate, (N=18) |  |  | simultaneous<sub>varied-f</sub>, (N=16) |  |  |
| --- | --- | --- | --- | --- | --- | --- | --- | --- | --- |
|  | X(mm) | Y(mm) | Z(mm) | X(mm) | Y(mm) | Z(mm) | X(mm) | Y(mm) | Z(mm) |
| Index | -44.0±1.0 | -24.1±1.7 | 57.3±1.3 | -47.6±1.1 | -23.5±1.8 | 54.5±0.4 | -47.7±1.6 | -19.7±2.0 | 52.7±1.5 |
| Middle | -44.0±1.0 | -24.1±1.7 | 57.2±1.3 | -45.7±1.2 | -24.3±1.4 | 58.4±0.3 | -47.3±1.6 | -20.5±2.0 | 53.7±1.3 |
| Ring | -44.0±1.0 | -24.2±1.6 | 57.1±1.3 | -44.2±0.9 | -25.3±1.5 | 58.4±0.4 | -45.8±1.3 | -23.8±1.6 | 57.3±1.8 |
| Little | -44.0±1.0 | -24.2±1.6 | 57.3±1.3 | -43.4±1.2 | -25.9±1.5 | 56.6±0.4 | -48.5±1.4 | -23.1±2.0 | 54.9±2.0 |

**Figure 4.**
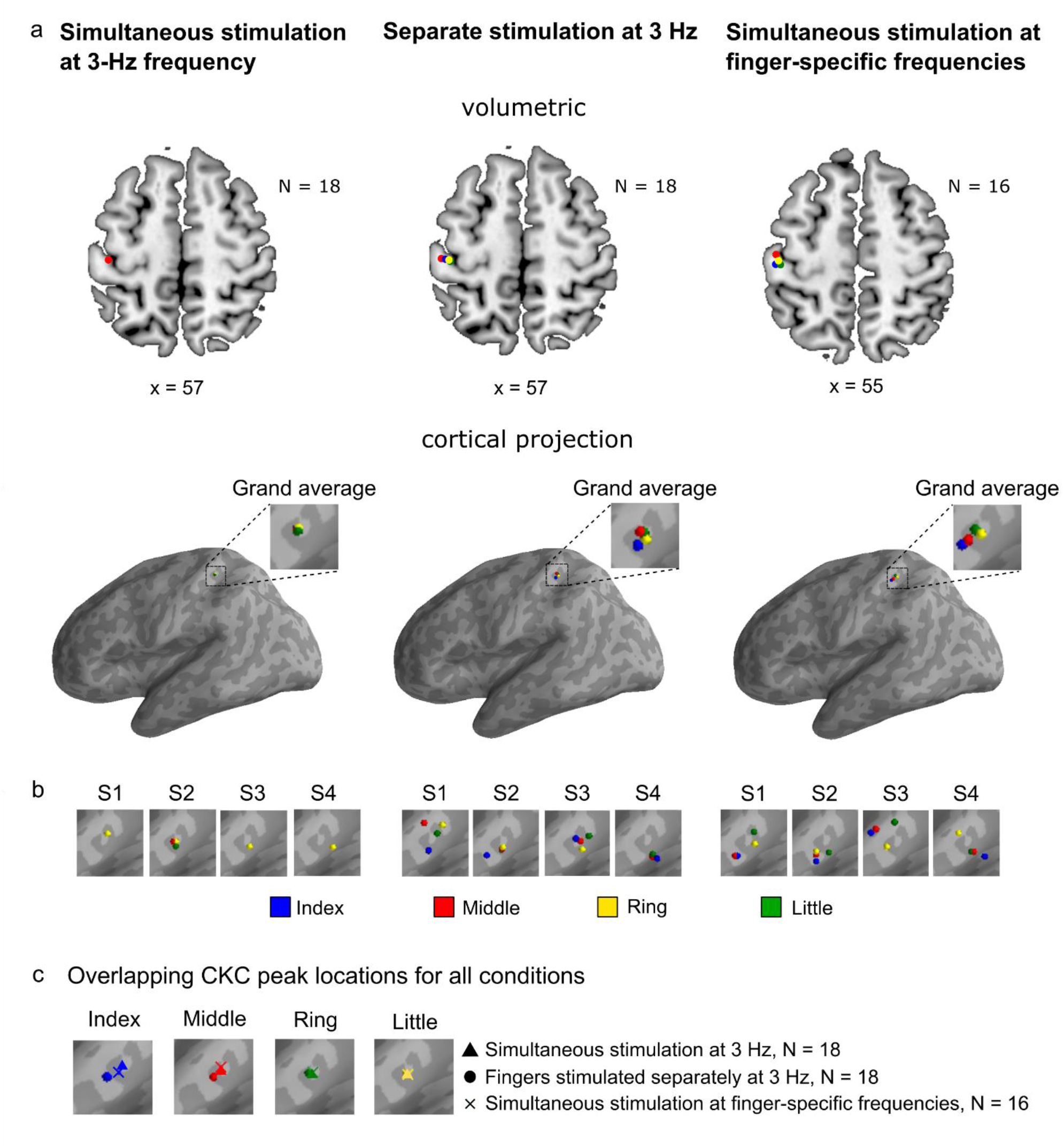
CKC-source-peak locations. (a) Group-level CKC source locations of each finger overlaid on the same volumetric brain (upper row) and cortical surface (lower row) separately for each condition. Please, note that x-coordinates are averages over the x-directional MNI coordinates of the CKC source locations of the four fingers (for the MNI source coordinates of each finger, see Table 3). (b) CKC source locations of each finger of four representative participants (S1–S4) overlaid on the same cortical surface separately for each condition. In S1 and S2 participant, the source locations of the index and little fingers roughly followed the somatotopic arrangement with respect to each other. The index finger was represented more dorsal and inferior to the little finger. (c) Group-level CKC source locations of each condition overlaid on the same cortical surface separately for each finger. Please, note that the source locations were concentrated on the Rolandic hand region of the SM1 cortex (*i*.*e*. central sulcus) in the source volume, but were misleadingly projected away from the central sulcus in the anterior wall of the postcentral sulcus when visualized to the cortical surface.

When stimulated simultaneously at the finger-specific frequencies, the CKC peak located 2.7 mm more medial for the ring finger than for the little finger (x = –48.5 ± 1.4 mm, p < 0.004). The CKC for the ring finger peaked 4.2 mm (p < 0.001) and for the little finger 3.5 mm (p < 0.001) more posterior than for the index finger (y = –19.6 ± 2.1 mm). Additionally, the CKC for the ring finger peaked 13.3 mm (p < 0.001) and for the little finger 12.6 mm (p < 0.001) more posterior than for the middle finger (y = –10.5 ± 1.8 mm). The CKC peak located 4.6 mm (p < 0.001) more superior for the ring finger and 2.2 mm (p < 0.05) more superior for the little finger than for the index finger (z = 52.7 ± 1.5 mm). Moreover, the CKC peaked 3.6 mm more superior for the ring finger than for the middle finger (z = 53.7 ± 1.3 mm p < 0.001) and 2.4 mm more superior for the ring finger than for the little finger (z = 54.9±2.0 mm, p < 0.006). No statistically significant differences were found between the CKC peak locations of any of the finger when stimulated separately *versus* simultaneously at the finger-specific frequencies (**Fig. 4c**).

## Discussion

We examined the CKC strength and cortical source location to proprioceptive stimulation of the right-hand fingers. Our results indicated that the *strongest* CKC was obtained with the most comprehensive stimulation of all four right-hand fingers *at the same 3-Hz frequency*. This approach resulted in about 64% stronger CKC than stimulation the fingers separately. CKC was weakest for the simultaneous stimulation of the fingers at finger-specific frequencies (2–3.5 Hz), being about 30% weaker than the CKC obtained with stimulation of the fingers separately and about 57% weaker than the strongest CKC. The CKC strength seems to be affected by both the finger dexterity and functional dominance. The CKC was weaker for dexterous fingers during separate stimulation in agreement with the neural efficiency hypothesis, but the opposite was true during simultaneous stimulation at finger-specific frequencies supporting the hypothesis of the functional dominance of the dexterous fingers in complex multidigit movements. All CKC source locations were concentrated (within ∼8 mm) in the Rolandic hand region of the SM1 cortex with some differences but without a consistent somatotopic order between the fingers. The simultaneous stimulation of all or several fingers can be suggested to improve robustness (signal-to-noise ratio) and time-efficiency of functional localization of the SM1 cortex of the hand when using the CKC method in combination with MEG. Finally, it remains open whether, in humans, the anatomical fractionation of the proprioceptive representations between the fingers of the same hand are less distinct and/or different compared to better studied tactile representations (Nakamura et al., 1998).

### Stronger CKC to simultaneous than separate-finger stimulation at 3 Hz

In agreement with our hypothesis, the sensor-level CKC was ∼64% stronger for the 3-Hz simultaneous simulation of the fingers (index, middle, ring and little) than for the 3-Hz stimulation of the fingers separately. The result was replicated in the source-level analysis which showed ∼28% stronger CKC for the simultaneous stimulation. Our result extends previous studies that have stimulated the proprioceptors related to the index finger (Bourguignon et al., 2016; Piitulainen et al., 2015, 2018b, 2020). We showed that the coherent proprioceptive afference from all induced fingers sum up to the SM1 cortex proprioceptive processing. Thus, the more comprehensive is the proprioceptive afference, the stronger is the cortical response or related proprioceptive processing. The stronger CKC may therefore reflect multidigit converging of the proprioceptive input. Similarly, the efferent motor output from the motor cortex converge and diverge when activating the hand muscles (McKiernan et al., 1998; Uematsu et al., 1992). Moreover, the fingers of the hand are functionally and anatomically overlapping. For example, activations of different neuromuscular regions in the monkey flexor digitorum profundus muscle, have shown to produce uniquely distributed tension in all five digits (Schieber et al., 2001). Thus, the neural control of the hand is likely more optimized for synergistic movements by combinations of fingers rather than control of individual fingers. Therefore, it is likely that the cortical proprioceptive processing is better optimized for the collective hand than individual digit movements. This hypothesis is further supported by a fMRI study which revealed that hand postural information, encoded through kinematic synergies of the fingers, strongly correlated with BOLD activation patterns in the SM1 cortex (Leo et al., 2016). An additional reason for the stronger CKC for the simultaneous finger stimulation could be the insufficient specificity of MEG to perfectly register the individual finger responses. MEG is biased towards neuronal activity from tangential currents, thus recording activity predominantly from sulci (*i*.*e*. fissural cortex) rather than gyri (Hillebrand and Barnes, 2002).

### Weaker CKC to simultaneous stimulation at finger-specific frequencies than separate stimulation

In contrast to our hypothesis, the stimulation of the fingers simultaneously at the finger-specific frequencies did not elicit analogous CKC values to their separate stimulation, but all fingers showed 30–36% weaker CKC when compared to the separate stimulation. Given that CKC has shown to be unaffected by the movement frequency of the finger (Marty et al., 2015; Piitulainen et al., 2015), it seems that proprioceptive afference from the other simultaneously stimulated fingers distracts the finger acceleration phase-locking to MEG signals reducing the CKC strength. As the fingers were stimulated with different frequencies, it is likely that the respective cortical responses are temporally overlapping in random manner, which likely hinders the respective signal-to-noise ratios and prominence of the MEG response (*i*.*e*., the coherent event), and eventually the CKC strength.

The greatest reduction in CKC from separate to simultaneous stimulation was observed for the little (∼48–62% weaker CKC) and ring (∼24–31%) fingers. This observation may be due to their lower level of dexterity and thus extent of the neuronal circuit responsible for the cortical proprioceptive processing for these fingers. It is plausible that during the simultaneous stimulation the MEG signal is more dominated by the more dexterous fingers, as the index and middle fingers, which may have larger cortical neuronal population involved in their proprioceptive processing. It is possible that cortical neural circuits of the more dexterous fingers partly overlap and dominate the circuits of the less dexterous (“assisting”) fingers. When a dominant finger is moved, proprioceptive afference (primarily from muscle afferents) would spread widely to the neural circuits of the other fingers, distracting the phase locking of their MEG responses. Ejaz at al. (2015) investigated fMRI data measured during active finger tapping tasks and showed that, indeed, BOLD representations of all fingers overlapped in the hand SM1 cortex, and the overlap was especially large between the middle and ring fingers. Furthermore, the between-finger similarity in BOLD response patterns correlated with the co-occurrence of common everyday-hand kinematics. This result suggested that the neural synergies are stronger between the fingers that frequently move together.

Another aspect is that the proprioceptive afference and its cortical processing may partly overlap in the functionally closely related fingers of the same hand. The functional interconnections between fingers have been estimated by measuring finger kinematics and kinetics when the subject has been instructed to produce isolated one finger contraction (Reilly and Hammond, 2000) or repetitive tapping (Aoki et al., 2003; Häger-Ross and Schieber, 2000). Strong “involuntary” forces or large movements by the noninstructed fingers were interpreted to reflect strong structural (*i*.*e*. tendons and muscles) and/or neuronal connection between the fingers. According to these studies, the index finger was the most independent and the ring finger the least independent of the other fingers. The middle finger was reported to be more independent than the little finger (Aoki et al., 2003; Reilly and Hammond, 2000) or vice versa (Häger-Ross and Schieber, 2000). Similar results have been obtained when the independence of the finger has been estimated based on the degree of how well the kinetics of the other fingers predict the finger kinematics in everyday-hand movements (Ingram et al., 2008). These results agree with our finding that the CKC strength of the most independent index finger was least affected by the simultaneous movement of the other fingers. Finally, based on our results, it appears that the level of independence and functional overlap in the fingers kinematics and functions are evident also in the cortical level of proprioceptive processing.

### CKC strength reflects finger dexterity and functional dominance

The dexterity (Kinoshita et al., 1996; Swanson et al., 1974; Zatsiorsky et al., 1998) and independence (Aoki et al., 2003; Häger-Ross and Schieber, 2000; Ingram et al., 2008; Reilly and Hammond, 2000) varies between the fingers based on the kinetics of voluntary finger actions. For this reason, we expected that the most dexterous index finger, with presumably the greatest degree of proprioceptive afference to the SM1 cortex, would show the strongest CKC, and the opposite would be true for the less dexterous fingers. Interestingly, we observed stronger CKC for the little and ring fingers (by ∼14–26%) than the middle and index fingers. This suggests that the stronger CKC may reflect weaker motor performance (*i*.*e*. dexterity) and/or level of usage of the ring and little fingers. The index and middle fingers are more utilized *e*.*g*. for grasping than the ring and little fingers (Kamakura et al., 1980). In addition, stronger CKC has shown to reflect worse standing balance performance in older (66–73 years) and younger (18–31 years) adults (Piitulainen et al., 2018b). Moreover, BOLD responses are stronger for the little than index finger tapping, presumably reflecting more challenging and/or less efficient cortical motor control of the less dexterous little finger (Erdler et al., 2001). Similarly, movement-related cortical potentials in the SM1 cortex have shown to be stronger for novices than motor-skilled subjects (Kita et al., 2001; Wright et al., 2012). Together, these results support the neural efficiency hypothesis, where a smaller neuronal population is recruited with improved motor efficiency and precision (Haier et al., 1988).

An opposite association was obtained when the fingers were stimulated at the finger-specific frequencies simultaneously. CKC was ∼43–77% stronger for the index, middle and ring fingers than for the little finger. In addition, the index and ring fingers yielded 35% stronger CKC than the middle finger. These results further demonstrate that the phase-locking of the MEG response and individual finger kinematics is affected by the other fingers at a finger-specific manner. It could be hypothesized that the more dexterous fingers dominate or “lead” the cortical proprioceptive processing during complex movement sequences of the hand. It is noteworthy that more statistically significant between-finger differences were detected at the source than sensor level analysis of the same data. This may reflect that the source analysis yields higher signal-to-noise ratio than the sensor analysis due to spatial filtering suppressing the irrelevant background activity. In addition, in the source space the contribution of all MEG sensors is taken into account when estimating CKC, whereas in sensor space only the one peak gradiometer pair contributes to the results. However, both approaches are acceptable as the main results were well replicated.

### CKC source locations were concentrated on the hand region of the SM1 cortex

Our fourth hypothesis was that the cortical source location would not significantly vary between the fingers in our participants, and thus each finger representation would similarly represent the Rolandic hand region in the SM1 cortex. In agreement with our hypothesis, the source locations of the fingers were only partly distinct and did not follow the consistent somatotopic pattern indicated by Penfield’s homunculus (Penfield and Boldery, 1937; Penfield and Rasmussen, 1950). However, there is no prior MEG evidence about proprioceptive representations of the same hand in the human SM1 cortex, although there is some evidence about somatotopic finger organization in cutaneous tactile domain (Nakamura et al., 1998).

As expected, the peak CKC locations were concentrated on the Rolandic SM1 cortex (within ∼8 mm) replicating previous results obtained by proprioceptive stimulation of the index finger (Bourguignon et al., 2016; Piitulainen et al., 2020, 2018b, 2015). However, the exact spatial coordinates for CKC source have been reported previously only for passive index finger movements elicited by an experimenter (Piitulainen et al., 2013b), not by precise stimulator. The distance between their and the current mean CKC source locations was ∼10 mm.

MEG is biased towards neuronal activity in the sulci (i.e., fissural cortex) and is less sensitive to deep and radial currents (Hamalainen et al., 1993; Hillebrand and Barnes, 2002). It is possible that due to these methodological limitations of MEG, we were unable to define the consistent proprioceptive finger representations in the SM1 cortex. Alternatively, the result may reflect that true neuroanatomy is less fractionated in the proprioceptive domain and could thus vary more between individuals when compared to, *e*.*g*., tactile domain. This is a challenge when distinguishing the group-level finger representations. A recent fMRI study revealed that finger-specific BOLD activation patterns elicited by finger tapping in the SM1 cortex are not somatotopically organized, and that their spatial layout is variable across subjects, while the relative similarity between any pair of activity patterns (*i*.*e*., Mahalanobis distances between digit-specific activation patterns) is invariant across subjects (Ejaz et al., 2015).

Our results are in line with tactile MEG and EEG studies that have reported either overlapping locations of somatosensory evoked field potentials of fingers (Baumgartner et al., 1993; Kalogianni et al., 2018; Schaefer et al., 2002; Simões et al., 2001) or that have managed to discriminate the activations related to mainly the index finger and thumb (Barbati et al., 2006; Nierula et al., 2013; Rossini et al., 2001, 1998). However, some MEG studies have found somatotopic cortical organization to tactile stimulation of the fingers (Nakamura et al., 1998). Similarly, fMRI studies have found somatotopic but partly overlapping cortical organization in S1 cortex to tactile stimuli of the fingers (Besle et al., 2013; Martuzzi et al., 2014).

### Further perspectives and limitations

Our results have practical implications for the functional mapping of the hand area in the SM1 cortex using CKC. The strongest CKC was obtained for the simultaneous 3-Hz stimulation of the fingers. Therefore, we suggest the *simultaneous* stimulation of several fingers at the *same* frequency to further improve robustness and time-efficiency of CKC method for functional mapping of the hand region in the SM1 cortex. However, this is not crucial issue as strong and robust CKC is detected for one-finger stimulation as well (Piitulainen et al., 2015), even in presence of strong magnetic artefacts (Bourguignon et al., 2016).

If the CKC strength of the individual fingers is of interest, each of the fingers should be stimulated separately rather than simultaneously at the finger-specific frequencies. This is because the simultaneous stimulation at the finger-specific frequencies resulted in the weaker CKC and, therefore, the signal-to-noise ratio likely decreases compared to the stimulation of the fingers separately.

Moreover, since the reduction in the CKC strength was finger-specific, the simultaneous stimulation with finger-specific frequencies can be less reliable approach to investigate the relative extent of the fingers proprioceptive processing in the SM1 cortex.

The proprioceptive stimulation of the fingers were generated with our neuroimaging compatible four-finger movement actuator which is an extension of the previous one-finger movement actuator (Piitulainen et al., 2015). The actuator had a millisecond timing accuracy and stabile stimuli and did not produce any artifacts to MEG signals. Thus, it provides a robust and reliable neuroimaging compatible tool to locate and investigate multi-finger proprioceptive afference to the SM1 cortex. The stimulator is suitable to study mechanisms of various motor disorders, since it allows meaningful reproducible comparisons between controls and patients who might have impaired ability to perform active motor tasks. However, it should be noted that the proprioceptive processing in the SM1 cortex may differ between passive and active movements and therefore, the four-finger actuator can only be used to investigate the processing of passive component of proprioception. Passive movements together with motor imaginary would correspond more closely the active movements, and may also be beneficial in the rehabilitation of the neurological patients. Indeed, imagined movements have shown to engage the same sensorimotor mechanisms as active movements do (Kilteni et al., 2018; Miller et al., 2010; Szameitat et al., 2006). Finally, the passive movement actuator does not activate solely the proprioceptors but inevitably also the functionally closely related tactile mechanoreceptors of the skin. These mechanoreceptors, responding, *e*.*g*., to stretch of the skin, can therefore be considered as a part of the same system providing the brain relevant information about the peripheral movement and actions. In addition, the CKC strength has shown to be unaffected by the level of tactile stimulation of the fingertip during active and passive index-finger movements (Piitulainen et al., 2013b), and therefore, CKC primarily reflects cortical processing of proprioceptive afference.

## Conclusion

The most comprehensive *simultaneous* stimulation of the right-hand fingers at the *same* frequency elicited the *strongest* CKC and can, therefore, be recommended as a robust and fast method for functional localization of the human hand region in the SM1 cortex using MEG. The modulation of the CKC strength in an individual finger by the other simultaneously stimulated fingers suggest that the respective proprioceptive afference is being processed in partly overlapping cortical neuronal circuits or populations. Individual fingers CKC strength was stronger in less dexterous or independent fingers in accordance with the neural efficiency hypothesis, but opposite observation was true when the fingers were stimulated simultaneously, which underlines the dominance of the more dexterous fingers in the cortical proprioceptive processing. The CKC sources of the fingers were concentrated in the Rolandic hand region of the SM1 cortex without systematic somatotopic organization, and thus, their representations appear partly overlapping, and/or MEG method is not sufficient to separate proprioceptive finger representations of the same hand adequately.

## Materials and methods

### Participants

Twenty-one healthy participants (mean age: 27.8, SD: 4.9, range: 20–40, 10 females, mean handedness score: 77.1; SD: 41.3 range: –80–100, one left-handed, one ambidextrous) without neuropsychiatric diseases, movement disorders or non-removable metallic objects in their body volunteered in the study. The data of three participants were excluded from the comparisons of CKC strength or source locations between *simultaneous* _constant-*f*_ and *separated* conditions and five between *simultaneous* _varied-*f*_ and *separated* conditions because of bad signal quality. Thus, the total numbers of participants included to the final analyses were 18 (mean age: 27.5, SD: 5.2, range: 20–40, 8 females, mean handedness score: 75.2; SD: 44.1 range: –80–100, one left-handed, one ambidextrous) and 16 (mean age: 27.2, SD: 5.4, range: 20–40, 7 females, mean handedness score: 73.6; SD: 46.6 range: –80–100, one left-handed, one ambidextrous), respectively. The handedness scores were assessed by a modified Edinburgh Handedness Inventory (Oldfield, 1971). The study was approved by the ethics committee of Aalto University, and the participants gave written informed consent before participation.

### Experimental design

At the beginning of the MEG session, the participant was briefed about the experiment. Before entering the MEG, the participant was provided with non-magnetic clothes and asked to remove any metallic objects he/she was wearing. During the MEG measurement, the participant was sitting with stimulated right hand on the custom-made proprioceptive stimulator (*i*.*e*. MEG-compatible movement actuator) placed on the table. The stimulator was an extension of our previously developed one-finger stimulator (Piitulainen et al., 2015). The tip of each of the four fingers was taped at the end of the finger-specific pneumatic muscle of the stimulator. Additionally, a piece of surgical tape (Leukoplast) was lightly attached on the palmar surface of each fingertip to minimize tactile stimulation elicited by the tactile contact between the fingertips and the stimulator. Accelerations of the fingers were measured with 3-axis accelerometers (ADXL335 iMEMS Accelerometer, Analog Devices Inc., Norwood, MA, USA) firmly taped on the nail of each finger. The left hand was resting on the thigh. The participant wore earplugs and Brownian noise was played in the background via a flat-panel speaker (Panphonics 60 × 60 SSHP, Tampere, Finland) to minimize auditory noise resulting from the airflow within the pneumatic muscles. To prevent the participant from seeing the moving fingers, a white A3-sized paper sheet was taped vertically to the MEG gantry. The participant was presented with a video of different landscapes (for two participants the video was not presented because of technical problems). Proprioceptive stimuli were controlled using Presentation software (ver. 18.1, Neurobehavioral Systems, Albany, CA, United States).

There were three conditions: (1) simultaneous stimulation of all four fingers at 3 Hz (*i*.*e*. stimulus onset asynchrony of 333 ms) in three 1-min bursts (*simultaneous* _constant-*f*_, 3 min stimulation in total), (2) stimulation of each finger separately at 3 Hz in three 1-min bursts (*separate*, 3 min stimulation per finger in total) and (3) simultaneous stimulation at finger-specific frequencies (at 2, 2.5,3 and 3.5 Hz, *simultaneous* _varied-*f*_) for 4 min. The data for *simultaneous* _constant-*f*_ and *separate* was collected in the same measurement, and the presentation order of the stimulation bursts was randomized for each participant. The duration of this measurement was 15 min.

### Data acquisition

MEG recordings were performed in a three-layer µ-metal magnetically shielded room (Imedco AG, Hägendorf, Switzerland) at MEG core of Aalto Neuroimaging Infrastructure (ANI) using a whole-scalp MEG device (Vectorview 4-D Neuromag Oy, Finland), with 204 gradiometer and 102 magnetometer sensors. MEG signals were band-pass filtered at 0.1–330 Hz and sampled at 1 kHz. Eye blinks were detected from electro-oculography (EOG) signal using an electrode pair placed above and below the left eye. Five head-position indicator (HPI) coils were used to determine the position of the head with respect to the MEG sensors, and to record head position continuously during the MEG recording. Prior the MEG measurements, the locations of the HPI coils were recorded with respect to three anatomical landmarks (nasion and two preauricular points) using a 3-D digitizer (Isotrack, Polhemus, Colchester, VT, USA). Additionally, points on the scalp surface (∼100) were digitized to facilitate co-registration between MEG data and anatomical magnetic resonance images. The participants were measured in seated position and instructed to avoid blinking and remain stationary during the measurement. The acceleration signals measured by the accelerometers attached on the nail of each finger were low-pass filtered at 330 Hz and sampled at 1 kHz time-locked to MEG signals.

Anatomical magnetic resonance imaging (MRI) images were acquired using a 3-tesla MRI scanner (MAGNETOM Skyra, Siemens Healthcare, Erlangen, Germany) and a 32-channel receiving head coli at the Advanced Magnetic Imaging (AMI) centre of Aalto University. MRI data was measured with a high-resolution T1-weighted Magnetization Prepared Rapid Gradient Echo (MPRAGE) pulse sequence (repetition time (TR) = 2530 ms, echo time (TE) = 3.3 ms, flip angle = 7, 256 x 256 matrix, 176 sagittal slices, 1-mm resolution).

### MEG preprocessing

MEG data was first visually inspected to identify noisy channels. Next, the uncorrelated sensor noise was reduced using the oversampled temporal projection (OTP, Larson and Taulu, 2018) algorithm. The temporally extended signal space separation algorithm (tSSS, MaxFilter 2.2 software, Elekta Neuromag Oy, Helsinki, Finland (Taulu and Simola, 2006), buffer length: 16 sec, correlation limit: 0.95) was applied to the MEG data to reduce environmental magnetic noise and interpolate the noisy channels. Visually identified noisy channels were given as an argument to the OTP and tSSS algorithms, and an automatic noisy channel detection (autobad option) was used in tSSS to further identify any noisy channels. To remove eye blinks and heart beats from the MEG signals, the data was decomposed into 30 independent components using fast independent component analysis (FastICA, Hyvärinen and Oja, 2000). Independent components related to the blinks and heart beats were identified by visually inspecting the topographies and time-series of ICA components and, thereafter, subtracted from the data. The ICA components were determined from the data filtered between 1–40 Hz using a zero-phase finite impulse response filter (firwin in SciPy 1.2.1; Hamming window, Virtanen et al., 2020) and removed from the nonfiltered data. OTP and ICA were performed using MNE Python software (version 3.6, Gramfort et al., 2014, 2013). The acceleration data of four accepted participants was missing and, therefore, replaced with the accelerometer data from another participant (stimulus sequence was identical across participants).

### Sensor level CKC analysis

To compute CKC between MEG and accelerometer signals for each finger, continuous data were split into epochs of 2000 ms with an overlap of 5 ms (Bortel and Sovka, 2007; Bourguignon et al., 2011). The epochs exceeding 2000 fT/cm at gradiometers and 4000 fT at magnetometers in peak-to-peak amplitude were excluded automatically from the data. Acceleration corresponding to each epoch was computed as Euclidean norm of the tree orthogonal accelerometer signals band-passed between 0.5– 195 Hz. The acceleration epochs were normalized by their Euclidean norms (Bourguignon et al., 2011). Thereafter, CKC was computed between MEG and accelerometer signals resulting in cross-, power, and coherence spectra and cross-correlogram (Halliday et al., 1995). Peak CKC strength was determined as the maximum coherence at the stimulation frequency over all MEG channels for each participant. The topographic distributions of CKC were visualized using Fieldtrip software (Oostenveld et al., 2011). The threshold for statistical significance corrected for multiple comparisons was computed as follows separately for all fingers and conditions for each participant with the following equation

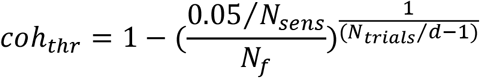

where *N*_*sens*_ is the number of MEG sensors among which the maximum coherence was searched, *N*_*f*_ is the number of the frequencies of interest (*i*.*e*. one since we studies only the movement frequency), *N _trials_* number of trials and *d* the overlap between trials (*i*.*e*. 5 ms, Bortel and Sovka, 2007).

### Source level CKC analysis

The dynamic imaging of coherent sources (DICS) beamformer (Bourguignon et al., 2013a, 2011; Gross et al., 2001) was used to estimate CKC between MEG signals and Euclidian norm of the accelerometer signals in the source space. To this end, cortical surfaces were reconstructed from T1 images using FreeSurfer’s recon-all algorithm (Freesurfer software v. 6.0, Dale et al., 1999; Fischl et al., 1999). To compute the forward model, a single-compartment boundary-element model (BEM) of the inner scull was generated using the FreeSurfer’s watershed algorithm. Each participant’s MEG sensor positions and MRI data were co-registered by aligning fiducial points in MEG and MRI (*i*.*e*. nasion, left and right preauricular points) as well as aligning MEG head digitization with the scalp. The fiducial points were manually identified on the MRI, and the fiducial registration error between MEG and MRI points was minimized by translating and rotating the MEG-digitized fiducials first automatically, and thereafter, adjusting the alignment manually. The forward model was computed for the volume source space with 6.2-mm spacing between the grid points (ico4 resolution). The leadfield with three components was reduced to the leadfield with two components corresponding to the highest singular values. The noise covariance matrix was estimated from the same file for which the source space CKC was computed. Finally, CKC maps were generated at the stimulation frequencies by computing CKC for all sources using DICS approach.

### Statistical analyses

First, we investigated whether the strength of CKC differs between the simultaneous 3-Hz stimulation and separate 3-Hz stimulation (H1). To this end, a two-way 2 x 4 repeated measurements analysis of variances (rANOVA) was carried out, with the within-participant factors of condition (*simultaneously* _*constant*_ vs. *separately*) and finger (index, middle, ring, little). Second, a similar rANOVA design was used to study whether CKC strength differs between the simultaneous stimulation at the finger-specific frequencies and separate 3-Hz stimulation (H2). For each finger, the number of accepted trials in separate stimulation was set as an upper limit of the trials in simultaneous stimulation at the finger-specific frequencies since the MEG measurement under separate stimulation was 1 minute shorter. We performed all rANOVAs separately for sensor and source level CKC strengths. Third, we used Newman-Keuls post hoc test to determine whether there were differences between fingers (H3). Fourth, we studied whether the source location of CKC differs between fingers in separate stimulation and simultaneous stimulation at the finger-specific frequencies by using Newman-Keuls post hoc comparisons separately for x-, y- and z-source coordinates in MNI space (H4). For all rANOVAs, Kolmogorov-Smirnov test and Mauchly test were run to test the normality and sphericity of the data, respectively. rANOVAs, Kolmogorov-Smirnov tests and Mauchly tests were implemented with Statistica 7.1 (StatSoft. Inc. 1984-2005). To test the consistency of the CKC location, we used the two-sample T-test (MATLAB R2019b) to compare the MNI coordinates of the CKC peaks elicited by *simultaneous* _constant-*f*_ condition and 3-Hz stimulation of the index finger. The normality of the data was tested for T-tests using Kolmogorov-Smirnov test (MATLAB R2019b, Massey, 1951).

## Acknowledgements

We thank Helge Kainulainen from his technical support in building the pneumatic system at Aalto NeuroImaging, Aalto University, Espoo, Finland. The reconstruction of cortical surfaces from T1 images was performed using computer resources within the Aalto University School of Science “Science-IT” project.

## Funding

This study has been supported by the Academy of Finland (grants #296240, #326988, #307250 and #327288) to HP and “Brain changes across the life-span” profiling funding (#311877) to University of Jyväskylä), and Jane and Aatos Erkko Foundation to HP.

## Author contributions

Hakonen, M.: Conceptualization, Methodology, Software, Validation, Formal Analysis, Data Curation, Writing – Original Draft Preparation, Visualization; Nurmi, T.: Investigation, Resources, Writing – Review & Editing; Jaatela, J.: Methodology, Writing – Review & Editing; Vallinoja, J.: Methodology, Writing – Review & Editing; Piitulainen H.: Conceptualization, Methodology, Writing – Review & Editing, Supervision, Project Administration, Funding Acquisition

## Notes

### Competing Interest Statement

The authors have declared no competing interest.

## References

Aoki, T., Francis, P.R., Kinoshita, H., 2003. Differences in the abilities of individual fingers during the performance of fast, repetitive tapping movements. Exp. Brain Res. 152, 270–280. https://doi.org/10.1007/s00221-003-1552-z

Barbati, G., Sigismondi, R., Zappasodi, F., Porcaro, C., Graziadio, S., Valente, G., Balsi, M., Rossini, P.M., Tecchio, F., 2006. Functional source separation from magnetoencephalographic signals. Hum. Brain Mapp. 27, 925–934. https://doi.org/10.1002/hbm.20232

Baumgartner, C., Doppelbauer, A., Sutherling, W.W., Lindinger, G., Levesque, M.F., Aull, S., Zeitlhofer, J., Deecke, L., 1993. Somatotopy of human hand somatosensory cortex as studied in scalp EEG. Electroencephalogr. Clin. Neurophysiol. Evoked Potentials 88, 271–279. https://doi.org/10.1016/0168-5597(93)90051-P

Besle, J., Sánchez-Panchuelo, R.M., Bowtell, R., Francis, S., Schluppeck, D., 2013. Single-subject fMRI mapping at 7 T of the representation of fingertips in S1: A comparison of event-related and phase-encoding designs. J. Neurophysiol. 109, 2293–2305. https://doi.org/10.1152/jn.00499.2012

Bortel, R., Sovka, P., 2007. Approximation of statistical distribution of magnitude squared coherence estimated with segment overlapping. Signal Processing 87, 1100–1117. https://doi.org/10.1016/j.sigpro.2006.10.003

Bourguignon, M., De Tiège, X., De Beeck, M.O., Ligot, N., Paquier, P., Van Bogaert, P., Goldman, S., Hari, R., Jousmäki, V., 2013a. The pace of prosodic phrasing couples the listener’s cortex to the reader’s voice. Hum. Brain Mapp. 34, 314–326. https://doi.org/10.1002/hbm.21442

Bourguignon, M., De Tiège, X., de Beeck, M.O., Pirotte, B., Van Bogaert, P., Goldman, S., Hari, R., Jousmäki, V., 2011. Functional motor-cortex mapping using corticokinematic coherence. Neuroimage 55, 1475–1479. https://doi.org/10.1016/j.neuroimage.2011.01.031

Bourguignon, M., Jousmäki, V., Marty, B., Wens, V., Op de Beeck, M., Van Bogaert, P., Nouali, M., Metens, T., Lubicz, B., Lefranc, F., Bruneau, M., De Witte, O., Goldman, S., De Tiège, X., 2013b. Comprehensive Functional Mapping Scheme for Non-Invasive Primary Sensorimotor Cortex Mapping. Brain Topogr. 26, 511–523. https://doi.org/10.1007/s10548-012-0271-9

Bourguignon, M., Piitulainen, H., De Tiège, X., Jousmäki, V., Hari, R., 2015. Corticokinematic coherence mainly reflects movement-induced proprioceptive feedback. Neuroimage 106, 382– 390. https://doi.org/10.1016/j.neuroimage.2014.11.026

Bourguignon, M., Whitmarsh, S., Piitulainen, H., Hari, R., Jousmäki, V., Lundqvist, D., 2016. Reliable recording and analysis of MEG-based corticokinematic coherence in the presence of strong magnetic artifacts. Clin. Neurophysiol. 127, 1460–1469. https://doi.org/10.1016/J.CLINPH.2015.07.030

Dale, A.M., Fischl, B., Sereno, M.I., 1999. Cortical surface-based analysis: I. Segmentation and surface reconstruction. Neuroimage 9, 179–194. https://doi.org/10.1006/nimg.1998.0395

Ejaz, N., Hamada, M., Diedrichsen, J., 2015. Hand use predicts the structure of representations in sensorimotor cortex. Nat. Neurosci. 18, 1034–1040. https://doi.org/10.1038/nn.4038

Erdler, M., Windischberger, C., Lanzenberger, R., Edward, V., Gartus, A., Deecke, L., Beisteiner, R., 2001. Dissociation of supplementary motor area and primary motor cortex in human subjects when comparing index and little finger movements with functional magnetic resonance imaging. Neurosci. Lett. 313, 5–8. https://doi.org/10.1016/S0304-3940(01)02167-X

Fischl, B., Sereno, M.I., Tootell, R.B., Dale, A.M., 1999. High-resolution intersubject averaging and a coordinate system for the cortical surface. Hum. Brain Mapp. 8, 272–84.

Gramfort, A., Luessi, M., Larson, E., Engemann, D.A., Strohmeier, D., Brodbeck, C., Goj, R., Jas, M., Brooks, T., Parkkonen, L., Hämäläinen, M., 2013. MEG and EEG data analysis with MNE-Python. Front. Neurosci. 7, 267. https://doi.org/10.3389/fnins.2013.00267

Gramfort, A., Luessi, M., Larson, E., Engemann, D.A., Strohmeier, D., Brodbeck, C., Parkkonen, L., Hämäläinen, M.S., 2014. MNE software for processing MEG and EEG data. Neuroimage 86, 446–460. https://doi.org/10.1016/j.neuroimage.2013.10.027

Gross, J., Kujala, J., Hämäläinen, M., Timmermann, L., Schnitzler, A., Salmelin, R., 2001. Dynamic imaging of coherent sources: Studying neural interactions in the human brain. Proc. Natl. Acad. Sci. U. S. A. 98, 694–699. https://doi.org/10.1073/pnas.98.2.694

Häger-Ross, C., Schieber, M.H., 2000. Quantifying the Independence of Human Finger Movements: Comparisons of Digits, Hands, and Movement Frequencies.

Haier, R.J., Siegel, B. V., Nuechterlein, K.H., Hazlett, E., Wu, J.C., Paek, J., Browning, H.L., Buchsbaum, M.S., 1988. Cortical glucose metabolic rate correlates of abstract reasoning and attention studied with positron emission tomography. Intelligence 12, 199–217. https://doi.org/10.1016/0160-2896(88)90016-5

Halliday, D.M., Rosenberg, J.R., Amjad, A.M., Breeze, P., Conway, B.A., Farmer, S.F., 1995. A framework for the analysis of mixed time series/point process data-Theory and application to the study of physiological tremor, single motor unit discharges and electromyograms. Prog. Biophys. Mol. Biol. https://doi.org/10.1016/S0079-6107(96)00009-0

Hamalainen, M., Hari, R., Ilmoniemi, R.J., Knuutila, J., Lounasmaa, O. V, 1993. Magnetoencephalography theory, instrumentation, and applications to noninvasive studies of the working human brain. https://doi.org/10.1103/revmodphys.65.413

Hari, R., Karhu, J., Hämäläinen, M., Knuutila, J., Salonen, O., Sams, M., Vilkman, V., 1993. Functional Organization of the Human First and Second Somatosensory Cortices: a Neuromagnetic Study. Eur. J. Neurosci. 5, 724–734. https://doi.org/10.1111/j.1460-9568.1993.tb00536.x

Hillebrand, A., Barnes, G.R., 2002. A quantitative assessment of the sensitivity of whole-head MEG to activity in the adult human cortex. Neuroimage 16, 638–650. https://doi.org/10.1006/nimg.2002.1102

Hyvärinen, A., Oja, E., 2000. Independent Component Analysis: algorithms and applications. Neural Netw 13, 411–430. https://doi.org/10.1016/S0893-6080(00)00026-5

Ingram, J.N., Körding, K.P., Howard, I.S., Wolpert, D.M., 2008. The statistics of natural hand movements. Exp. Brain Res. 188, 223–236. https://doi.org/10.1007/s00221-008-1355-3

Jerbi, K., Lachaux, J.P., N’Diaye, K., Pantazis, D., Leahy, R.M., Garnero, L., Baillet, S., 2007. Coherent neural representation of hand speed in humans revealed by MEG imaging. Proc. Natl. Acad. Sci. U. S. A. 104, 7676–7681. https://doi.org/10.1073/pnas.0609632104

Kalogianni, K., Daffertshofer, A., van der Helm, F.C.T., Schouten, A.C., de Munck, J.C., 2018. Disentangling Somatosensory Evoked Potentials of the Fingers: Limitations and Clinical Potential. Brain Topogr. 31, 498–512. https://doi.org/10.1007/s10548-017-0617-4

Kamakura, N., Matsuo, M., Ishii, H., Mitsuboshi, F., Miura, Y., 1980. Patterns of static prehension in normal hands. Am. J. Occup. Ther. 34, 437–445. https://doi.org/10.5014/ajot.34.7.437

Kilteni, K., Andersson, B.J., Houborg, C., Ehrsson, H.H., 2018. Motor imagery involves predicting the sensory consequences of the imagined movement. Nat. Commun. 9. https://doi.org/10.1038/S41467-018-03989-0

Kinoshita, H., Murase, T., Bandou, T., 1996. Grip posture and forces during holding cylindrical objects with circular grips. Ergonomics 39, 1163–1176. https://doi.org/10.1080/00140139608964536

Kita, Y., Mori, A., Nara, M., 2001. Two types of movement-related cortical potentials preceding wrist extension in humans. Neuroreport 12, 2221–2225. https://doi.org/10.1097/00001756-200107200-00035

Larson, E., Taulu, S., 2018. Reducing Sensor Noise in MEG and EEG Recordings Using Oversampled Temporal Projection. IEEE Trans. Biomed. Eng. 65. https://doi.org/10.1109/TBME.2017.2734641

Leo, A., Handjaras, G., Bianchi, M., Marino, H., Gabiccini, M., Guidi, A., Scilingo, E.P., Pietrini, P., Bicchi, A., Santello, M., Ricciardi, E., 2016. A synergy-based hand control is encoded in human motor cortical areas. Elife 5. https://doi.org/10.7554/eLife.13420

Martuzzi, R., van der Zwaag, W., Farthouat, J., Gruetter, R., Blanke, O., 2014. Human finger somatotopy in areas 3b, 1, and 2: A 7T fMRI study using a natural stimulus. Hum. Brain Mapp. 35, 213–226. https://doi.org/10.1002/hbm.22172

Marty, B., Bourguignon, M., Op de Beeck, M., Wens, V., Goldman, S., Van Bogaert, P., Jousmäki, V., De Tiège, X., 2015. Effect of movement rate on corticokinematic coherence. Neurophysiol. Clin. 45, 469–474. https://doi.org/10.1016/j.neucli.2015.09.002

Marty, B., Naeije, G., Bourguignon, M., Wens, V., Jousmäki, V., Lynch, D.R., Gaetz, W., Goldman, S., Hari, R., Pandolfo, M., De Tiège, X., 2019. Evidence for genetically determined degeneration of proprioceptive tracts in Friedreich ataxia. Neurology 93, E116–E124. https://doi.org/10.1212/WNL.0000000000007750

Massey, F.J., 1951. The Kolmogorov-Smirnov Test for Goodness of Fit. J. Am. Stat. Assoc. 46, 68– 78. https://doi.org/10.1080/01621459.1951.10500769

McKiernan, B.J., Marcario, J.K., Karrer, J.H., Cheney, P.D., 1998. Corticomotoneuronal postspike effects in shoulder, elbow, wrist, digit, and intrinsic hand muscles during a reach and prehension task. J. Neurophysiol. 80, 1961–1980. https://doi.org/10.1152/jn.1998.80.4.1961

Miller, K.J., Schalk, G., Fetz, E.E., Den Nijs, M., Ojemann, J.G., Rao, R.P.N., 2010. Cortical activity during motor execution, motor imagery, and imagery-based online feedback. Proc. Natl. Acad. Sci. U. S. A. 107, 4430–4435. https://doi.org/10.1073/pnas.0913697107

Nakamura, A., Yamada, T., Goto, A., Kato, T., Ito, K., Abe, Y., Kachi, T., Kakigi, R., 1998. Somatosensory homunculus as drawn by MEG. Neuroimage 7, 377–386. https://doi.org/10.1006/nimg.1998.0332

Nierula, B., Hohlefeld, F.U., Curio, G., Nikulin, V. V., 2013. No somatotopy of sensorimotor alpha-oscillation responses to differential finger stimulation. Neuroimage 76, 294–303. https://doi.org/10.1016/j.neuroimage.2013.03.025

Oldfield, R.C., 1971. The assessment and analysis of handedness: The Edinburgh inventory. europsychologia 9, 97–113. https://doi.org/10.1016/0028-3932(71)90067-4

Oostenveld, R., Fries, P., Maris, E., Schoffelen, J.M., 2011. FieldTrip: Open source software for advanced analysis of MEG, EEG, and invasive electrophysiological data. Comput. Intell. Neurosci. 2011. https://doi.org/10.1155/2011/156869

Penfield, W., Boldery, E., 1937. Somatic motor and sensory representation in the cerebral cortex of man as studied by electrical stimulation. Brain 60, 389–443. https://doi.org/10.1093/brain/60.4.389

Penfield, W., Rasmussen, T., 1950. The Cerebral Cortex of Man: A Clinical Study of Localization of Function. J. Am. Med. Assoc. 144, 1412. https://doi.org/10.1001/jama.1950.02920160086033

Piitulainen, H., Bourguignon, M., De Tiège, X., Hari, R., Jousmäki, V., 2013a. Coherence between magnetoencephalography and hand-action-related acceleration, force, pressure, and electromyogram. Neuroimage 72, 83–90. https://doi.org/10.1016/j.neuroimage.2013.01.029

Piitulainen, H., Bourguignon, M., De Tiège, X., Hari, R., Jousmäki, V., 2013b. Corticokinematic coherence during active and passive finger movements. Neuroscience 238, 361–370. https://doi.org/10.1016/j.neuroscience.2013.02.002

Piitulainen, H., Bourguignon, M., Hari, R., Jousmäki, V., 2015. MEG-compatible pneumatic stimulator to elicit passive finger and toe movements. Neuroimage 112, 310–317. https://doi.org/10.1016/J.NEUROIMAGE.2015.03.006

Piitulainen, H., Illman, M., Laaksonen, K., Jousmäki, V., Forss, N., 2018a. Reproducibility of corticokinematic coherence. Neuroimage 179, 596–603. https://doi.org/10.1016/J.NEUROIMAGE.2018.06.078

Piitulainen, H., Illman, M.J., Jousmäki, V., Bourguignon, M., 2020. Feasibility and reproducibility of electroencephalography-based corticokinematic coherence. J. Neurophysiol.

Piitulainen, H., Seipäjärvi, S., Avela, J., Parviainen, T., Walker, S., 2018b. Cortical Proprioceptive Processing Is Altered by Aging. Front. Aging Neurosci. 10, 147. https://doi.org/10.3389/fnagi.2018.00147

Reilly, K.T., Hammond, G.R., 2000. Independence of force production by digits of the human hand. Neurosci. Lett. 290, 53–56. https://doi.org/10.1016/S0304-3940(00)01328-8

Rossini, P.M., Tecchio, F., Pizzella, V., Lupoi, D., Cassetta, E., Paqualetti, P., 2001. Interhemispheric differences of sensory hand areas after monohemispheric stroke: MEG/MRI integrative study. Neuroimage 14, 474–485. https://doi.org/10.1006/nimg.2000.0686

Rossini, P.M., Tecchio, F., Pizzella, V., Lupoi, D., Cassetta, E., Pasqualetti, P., Romani, G.L., Orlacchio, A., 1998. On the reorganization of sensory hand areas after mono-hemispheric lesion: A functional (MEG)/anatomical (MRI) integrarive study. Brain Res. 782, 153–166. https://doi.org/10.1016/S0006-8993(97)01274-2

Schaefer, M., Mühlnickel, W., Grüsser, S.M., Flor, H., 2002. Reproducibility and stability of neuroelectric source imaging in primary somatosensory cortex. Brain Topogr. 14, 179–189. https://doi.org/10.1023/A:1014598724094

Schieber, M.H., Gardinier, J., Liu, J., 2001. Tension Distribution to the Five Digits of the Hand by Neuromuscular Compartments in the Macaque Flexor Digitorum Profundus.

Simões, C., Mertens, M., Forss, N., Jousmäki, V., Lütkenhöner, B., Hari, R., 2001. Functional Overlap of Finger Representations in Human SI and SII Cortices. J. Neurophysiol. 86, 1661– 1665.

Smeds, E., Vanhatalo, S., Piitulainen, H., Bourguignon, M., Jousmäki, V., Hari, R., 2017. Corticokinematic coherence as a new marker for somatosensory afference in newborns. Clin. Neurophysiol. 128, 647–655. https://doi.org/10.1016/j.clinph.2017.01.006

Swanson, A.B., Matev, I.B., De Groot, G., 1974. The strength of the hand. Inter Clin. Inf. Bull. 13, 1–8. https://doi.org/10.1016/0003-6870(72)90119-6

Szameitat, A.J., Shen, S., Sterr, A., 2006. Motor imagery of complex everyday movements. An fMRI study. https://doi.org/10.1016/j.neuroimage.2006.09.033

Taulu, S., Simola, J., 2006. Spatiotemporal signal space separation method for rejecting nearby interference in MEG measurements. Phys. Med. Biol 51, 1–10. https://doi.org/10.1088/0031-9155/51/0/000

Uematsu, S., Lesser, R., Fisher, R.S., Gordon, B., Hara, K., Krauss, G.L., Vining, E.P., Webber, R.W., 1992. Motor and sensory cortex in humans: Topography studied with chronic subdural stimulation. Neurosurgery 31, 59–72. https://doi.org/10.1227/00006123-199207000-00009

Virtanen, P., Gommers, R., Oliphant, T.E., Haberland, M., Reddy, T., Cournapeau, D., Burovski, E., Peterson, P., Weckesser, W., Bright, J., van der Walt, S.J., Brett, M., Wilson, J., Millman, K.J., Mayorov, N., Nelson, A.R.J., Jones, E., Kern, R., Larson, E., Carey, C.J., Polat, İ., Feng, Y., Moore, E.W., VanderPlas, J., Laxalde, D., Perktold, J., Cimrman, R., Henriksen, I., Quintero, E.A., Harris, C.R., Archibald, A.M., Ribeiro, A.H., Pedregosa, F., van Mulbregt, P., Vijaykumar, A., Bardelli A. Pietro, Rothberg, A., Hilboll, A., Kloeckner, A., Scopatz, A., Lee, A., Rokem, A., Woods, C.N., Fulton, C., Masson, C., Häggström, C., Fitzgerald, C., Nicholson, D.A., Hagen, D.R., Pasechnik, D. V., Olivetti, E., Martin, E., Wieser, E., Silva, F., Lenders, F., Wilhelm, F., Young, G., Price, G.A., Ingold, G.L., Allen, G.E., Lee, G.R., Audren, H., Probst, I., Dietrich, J.P., Silterra, J., Webber, J.T., Slavič, J., Nothman, J., Buchner, J., Kulick, J., Schönberger, J.L., de Miranda Cardoso, J.V., Reimer, J., Harrington, J., Rodríguez, J.L.C., Nunez-Iglesias, J., Kuczynski, J., Tritz, K., Thoma, M., Newville, M., Kümmerer, M., Bolingbroke, M., Tartre, M., Pak, M., Smith, N.J., Nowaczyk, N., Shebanov, N., Pavlyk, O., Brodtkorb, P.A., Lee, P., McGibbon, R.T., Feldbauer, R., Lewis, S., Tygier, S., Sievert, S., Vigna, S., Peterson, S., More, S., Pudlik, T., Oshima, T., Pingel, T.J., Robitaille, T.P., Spura, T., Jones, T.R., Cera, T., Leslie, T., Zito, T., Krauss, T., Upadhyay, U., Halchenko, Y.O., Vázquez-Baeza, Y., 2020. SciPy 1.0: fundamental algorithms for scientific computing in Python. Nat. Methods 17, 261–272. https://doi.org/10.1038/s41592-019-0686-2

Wright, D.J., Holmes, P., Di Russo, F., Loporto, M., Smith, D., 2012. Reduced Motor Cortex Activity during Movement Preparation following a Period of Motor Skill Practice. PLoS One 7, e51886. https://doi.org/10.1371/journal.pone.0051886

Zatsiorsky, V.M., Li, Z.M., Latash, M.L., 1998. Coordinated force production in multi-finger tasks: Finger interaction and neural network modeling. Biol. Cybern. 79, 139–150. https://doi.org/10.1007/s004220050466

